# Target Binding and Sequence Prediction With LSTMs

**DOI:** 10.1101/504415

**Authors:** Michael Teti, Rachel St Clair, Mirjana Pavlovic, Elan Barenholtz, William Hahn

**Affiliations:** with the CCSBS and CEECS, Florida Atlantic University, Boca Raton, FL, United States 33431 USA

**Author notes:** Correspondance (RS).: 1+) (RS).

**Keywords:** LSTM, Proteins, Binding Prediction, Primary Sequence

## Abstract

Deep recurrent neural networks (DRNNs) have recently demonstrated strong performance in sequential data analysis, such as natural language processing. These capabilities make them a promising tool for inferential analysis of sequentially structured bioinformatics data as well. Here, we assessed the ability of Long Short-Term Memory (LSTM) networks, a class of DRNNs, to predict properties of proteins based on their primary structures. The proposed architecture is trained and tested on two different datasets to predict whether a given sequence falls into a certain class or not. The first dataset, directly imported from Uniprot, was used to train the network on whether a given protein contained or did not contain a conserved sequence (homeodomain), and the second dataset, derived by literature mining, was used to train a network on whether a given protein binds or doesn’t bind to Artemisinin, a drug typically used to treat malaria. In each case, the model was able to differentiate between the two different classes of sequences it was given with high accuracy, illustrating successful learning and generalization. Upon completion of training, an ROC curve was created using the homeodomain and artemisinin validation datasets. The AUC of these datasets was 0.80 and 0.87 respectively, further indicating the models’ effectiveness. Furthermore, using these trained models, it was possible to derive a protocol for sequence detection of homeodomain and binding motif, which are well-documented in literature, and a known Artemisinin binding site, respectively [1-3]. Along with these contributions, we developed a python API to directly connect to Uniprot data sourcing, train deep neural networks on this primary sequence data using TensorFlow, and uniquely visualize the results of this analysis. Such an approach has the potential to drastically increase accuracy and reduce computational time and, current major limitations in informatics, from inquiry to discovery in protein function research.

## Introduction

Protein primary sequence is the biological language a cell uses to construct and maintain structural and functional networks/components. Position-specific residues and motifs dictate function, and elucidating these compositional domains can aid in a variety of applications (ie. drug discovery and detection). As proteomic datasets grow exponentially, the need for efficient methods in collecting, sorting, and analyzing becomes an increasingly imperative aspect of biological research. Traditional methods, which involve multiple sequence alignments (MSA) can be computationally expensive. Kamena, in a 2009 review of MSA in the high-throughput era, highlights computational limitations as a major deficit of current approaches, calling for big-data processing [4]. Furthermore, many of these methods involve two dominating constraints, reliance upon evolutionary data and inaccurate alignment sequencing: emphasizing the need of simple pipelines in protein FASTA organization that can *accurately* handle exponentially emerging data quantities [5]. One set of methods with considerable promise to have emerged in the last decade is the application of machine learning algorithms to protein primary sequence analysis pipeline. MultiLoc, one of the earliest successful pioneers, used SVM based models that predicted subcellular localization from SWISSprot sequence inputs along with largely hand-annotated secondary features (ie. structure), a major drawback to efficient computation [6]. However, SVMs and other, more dated machine learning techniques have significant limitations in larger, more complex datasets. Recently, multi-layer, or ‘deep’ neural networks have led to remarkable advances in multiple big data domains such as computer vision and natural language processing.

Because proteins carry information in their sequence of amino acids, it is appropriate to apply ‘recurrent’ networks that leverage previous inputs when making an inference on a current input. A particularly successful RNN in natural language processing is the Long Short Term Memory, or LSTM (Fig. 1) [6,7]. A unique feature of the LSTM is the network’s ability to receive information in a time series while a hidden state cell - which acts similar to a long-term memory mechanism - allows the information to be selectively fed-forward or withheld based on learned weights of the model. This ability to variably consider selected information of previous input is one of the primary reasons LSTMs have been used for language processing tasks.

**Fig. 1.**
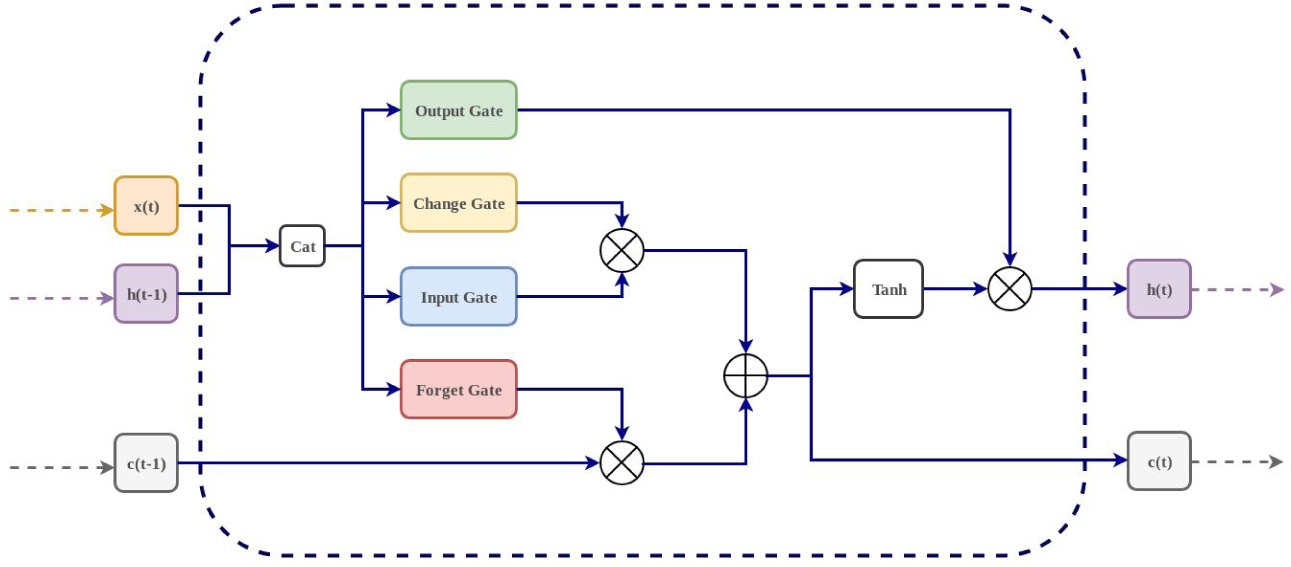
LSTM Node. This diagram shows the inner workings of each node in the LSTM used in this work.

**Fig. 2.**
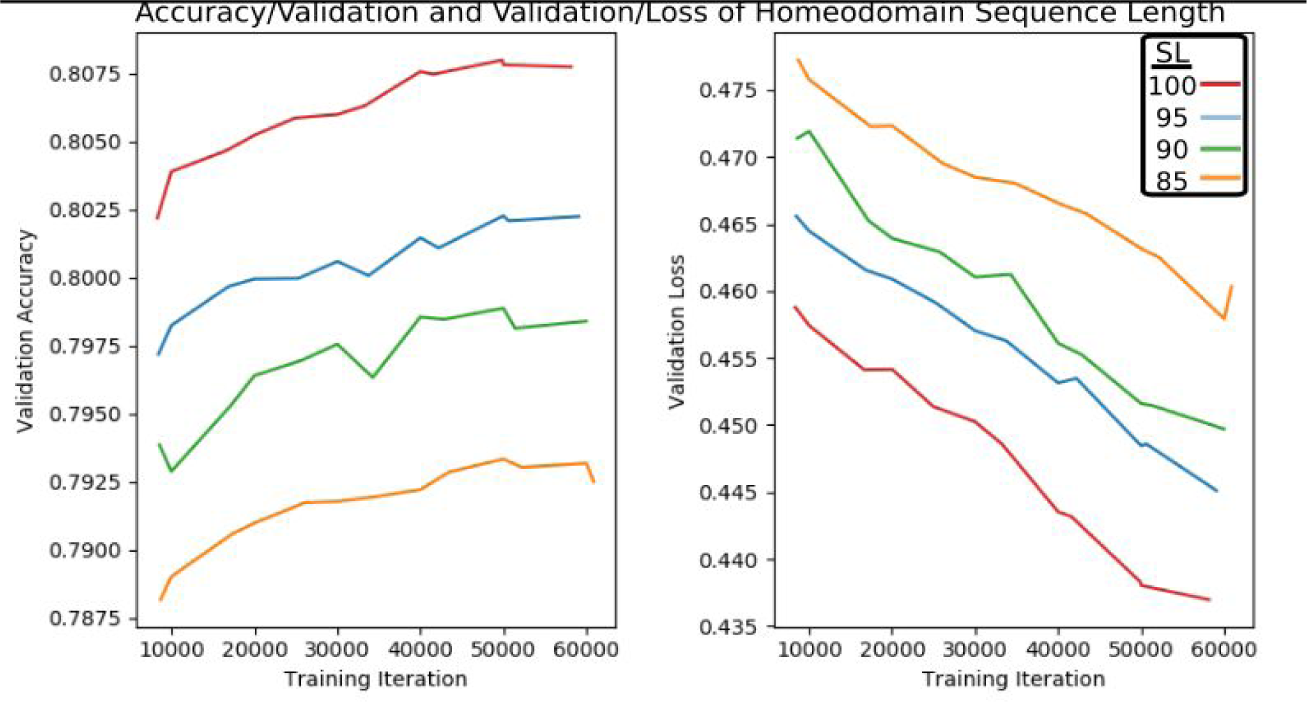
Homeodomain Accuracy. Calibration of LSTM model with homeoprotein datasets during phase one. Accuracy of prediction of classification was assessed at sequence lengths from 85-100 amino acids

**Fig. 3.**
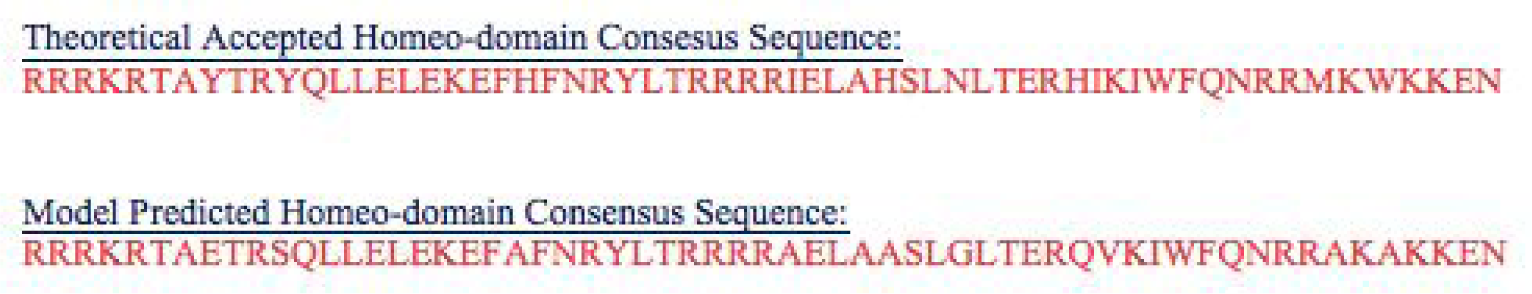
Theoretical vs. Predicted Homeodomain Consensus Sequence. The first sequence is the literature-accepted amino acid homeodomain consensus sequence. The second sequence is the predicted sequence the model showed most conserved among proteins containing the homeodomain sequence.

**Fig. 4.**
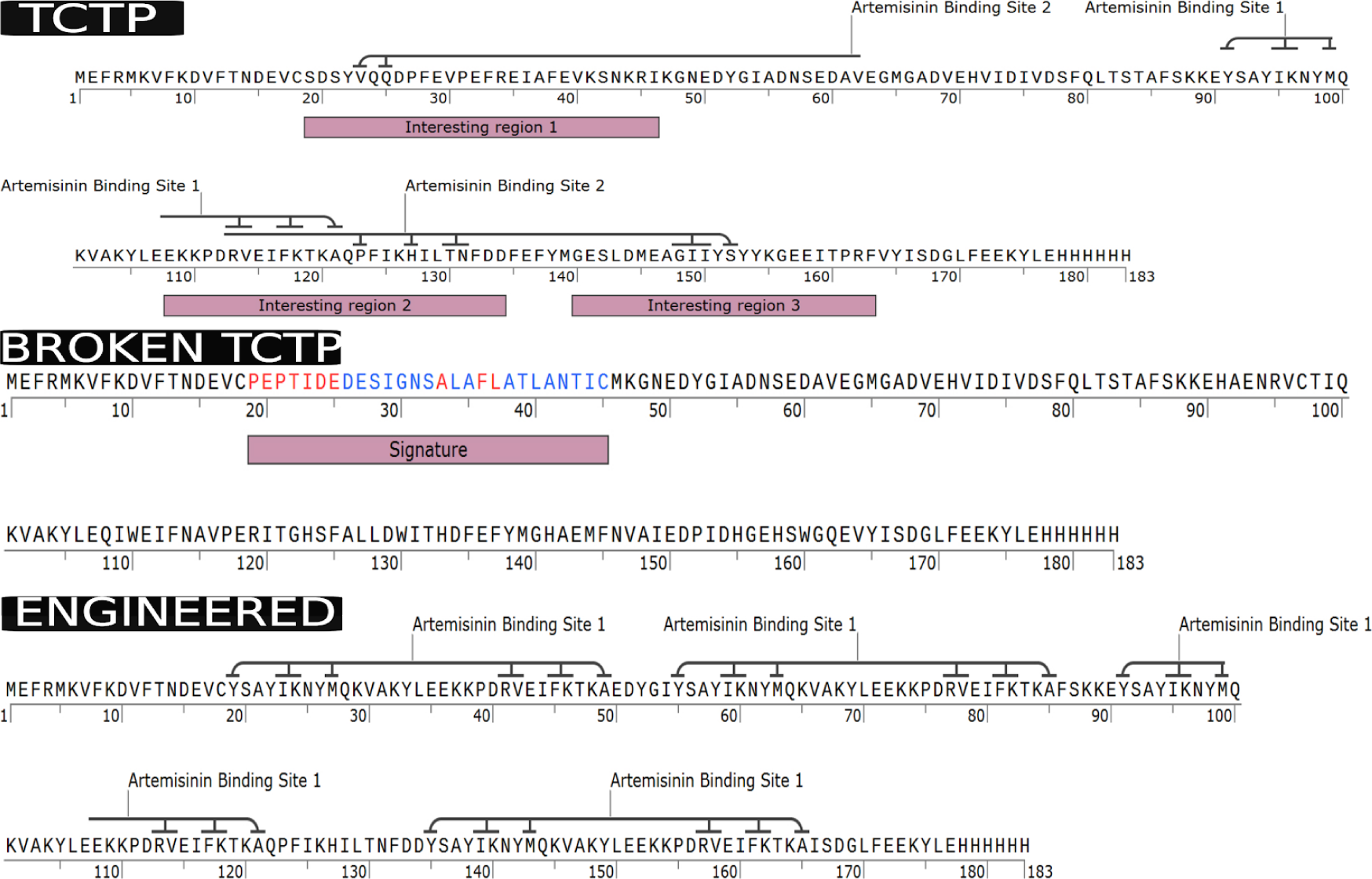
Three Artemisinin Proteins Used in Testing. These sequences are 1. accepted binding TCTP protein from literature 2. non-binding broken TCTP protein, 3. Theoretical genetically-engineered protein.

Due to protein primary sequences’ nature of position-specific residues dictating function, they are strong candidates for the application of LSTM networks as well. This potential is illustrated by a recent study which found that LSTM-based RNNs applied to protein sequences from UniProt were able to accurately predict several basic functions [8]. However, the computational methods employed in that study had several limitations. First, they only view proteins as a whole, a very computationally expensive task for longer proteins. Another drawback to the previous approach was using unbalanced validation datasets, possibly allowing for bias towards classifying proteins not in the functional category, rather than proteins in the specified category. Furthermore, the study removed proteins shorter than 10 residues and longer than 1000 to reduce false positives; in synthetic applications and some natural datasets, these outlier-sized proteins may be necessary.

In the current study we aimed to further validate as well as extend the application of LSTM networks in analyzing protein primary sequence data. We use padded sequence lengths and well as balanced validation datasets to ensure training and validation uniformity. In addition, we introduce a ‘sliding-window’ technique, described in the Methods section below, that is able to look at various parts of the protein in smaller chunks rather than processing each protein sequence as a whole, which becomes exponentially more computationally expensive as proteins increase in length. Finally, we introduce a method that goes beyond simple classification and aims to elucidate the compositional components (i.e. features) of the protein responsible for the model’s classification behavior by best-fit data inference from output layer nodes.

We first assessed whether LSTM models could determine whether given protein sequences contained a homeobox. The conservation of homeoboxes is widely regarded as a hallmark identifier in developmental proteins [3]. Since homeodomain is so widely conserved and is similar to a binding motif such as the one observed in Artemisinin-binding proteins, it served as a proof-of-concept for the models’ inference capabilities. Next, we attempted to extend the application of these models to a more challenging and potentially useful task: proteins that bind to antimalarial drug Artemisinin. Although some binding motifs such as translationally controlled tumour protein (TCTP) and ferritin are known to bind effectively, there are not extensive collections of Artemisinin binding proteins and properties [1,3,9]. Each motif resides in a protein’s primary structure, has position-specific residues, and similar architectural sequence components, yet may have residues that vary at unpredicted locations in the sequence. Classifying proteins that have such binding motif and elucidating what amino acids comprise that motif would be a step towards discovering essential binding properties necessary.

The proposed architecture provides this novel ability to predict binding motif in a cohort of amino acids by incorporating an end-to-end deep learning architecture: during model training, internal abstract representations of the functional component in the data-set’s class can be queried to find the recurrent domain. Next, by finding differences of each protein to the output layer nodes representing protein function, the most activated component is shown: feature selection. This technique utilizes learned features of protein class to predict the sequence creating the respective function. Since the features are learned without human discretion, the model is representing the binding motif by means of best-fit data inference. Sequence prediction thus transforms the basic classification model into a high precision suggestive tool for directly inferring functional amino-acid composition.

## Results

### Homeodomain Consensus Prediction

The model was first trained on the Homeodomain dataset with ‘sliding window’ technique to view each protein containing homeobox and not containing homeobox for classification task. Accuracy measures, shown in figure two, shows classification performance across four sequence lengths in this first Homeodomain task. The AUC measured 0.80 for sequence length 50 of such task (Fig. 8). As previously stated, the homeodomain dataset was used to validate the model’s ability to find a feature conserved amongst the dataset and show such feature’s amino acid composition. In the second task, prompted in the analysis phase for the sequence that most activated the output node that responds to homeodomain containing class with sequence length of 60 amino acids, the model returned a predicted sequence that only differed from the conserved homeobox literature-accepted sequence at specific residue positions by 10 amino acids, as seen in figure three. The sequence length of 60 was chosen due to the literature-accepted homeodomain being 60 amino acids long. [2] The variability in the predicted sequence is expected since a consensus sequence, even highly conserved, will vary at different positions in the sequence (the nature of degenerate biological code and variance in a population). This ability of the model to find homeodomain sequence in a dataset of homeodomain containing proteins, without being directly told what part of the proteins’ primary sequence is a homeodomain, is due to the models ability to extract the specific compositional component causing protein function (homeodomain activity). These results reinforce the models ability to analyze a dataset and determine important areas of interest within a certain class — in this case by analyzing a group of similar proteins compared to proteins not in the indicated group to predict consensus sequence.

### Artemisinin Binding Prediction

The model was then trained on Artemisinin-binding proteins with ‘sliding-window’ range of sequence lengths from 3-150 residues. After the model was trained, the network was given three novel sequences separate from the original dataset, shown in figure four. TCTP was used as a positive control sequence since literature validates its binding ability to Artemisinin [1]. A broken TCTP sequence with arbitrarily inserted amino acids to render it nonfunctional was used for the negative control since the known literature binding site was manually changed into an unrelated sequence. Finally the last sequence was hand-engineered to model the TCTP with other related functions a genetic engineer would recommend and need for wet-lab purposes. These three are the novel proteins given to the model to test its ability to classify binding or non-binding. These sequences were then used to test the model to confirm it could predict whether the TCTP known-to-bind molecule would bind and the broken TCTP would not bind, as well as determine if the human engineered sequence would bind or not. The model completed this task with good accuracy (Fig. 5). For protein sequences that were predicted by the model to bind, node one - the output layer in the network that predicted binding - had a higher confidence than node zero - which represented non-binding. The results (per test protein) are shown on the specified sequence length 50, thus creating a line graph corresponding each snippet of sequence to a prediction value (from 0-1) for both node one and node zero in figure five. For the broken TCTP protein, node zero was consistently higher than node one, indicating the model predicts this protein to be more likely non-binding. The TCTP protein was predicted to bind as indicated by node one consistently being more activated than node zero, indicating binding. The novel engineered sequence was predicted to bind with essentially no probability of non-binding (Fig. 5). Analysis phase determined which sequence fragment activated node one the most for a specified sequence length. For sequence length 50, the AUC was 0.87, showing the confidence of true positive for classification task (Fig. 9). Since the model is end-to-end (no human intervention in decision output), the conserved sequence the model found most preserved among the binding class could be determined. Although not all proteins in the Artemisinin binding dataset were TCTP analogs, it was noticed that this sequence predicted by the model is very similar to the TCTP proposed binding site shown in the red-highlighted regions of figure 6.

**Fig. 5.**
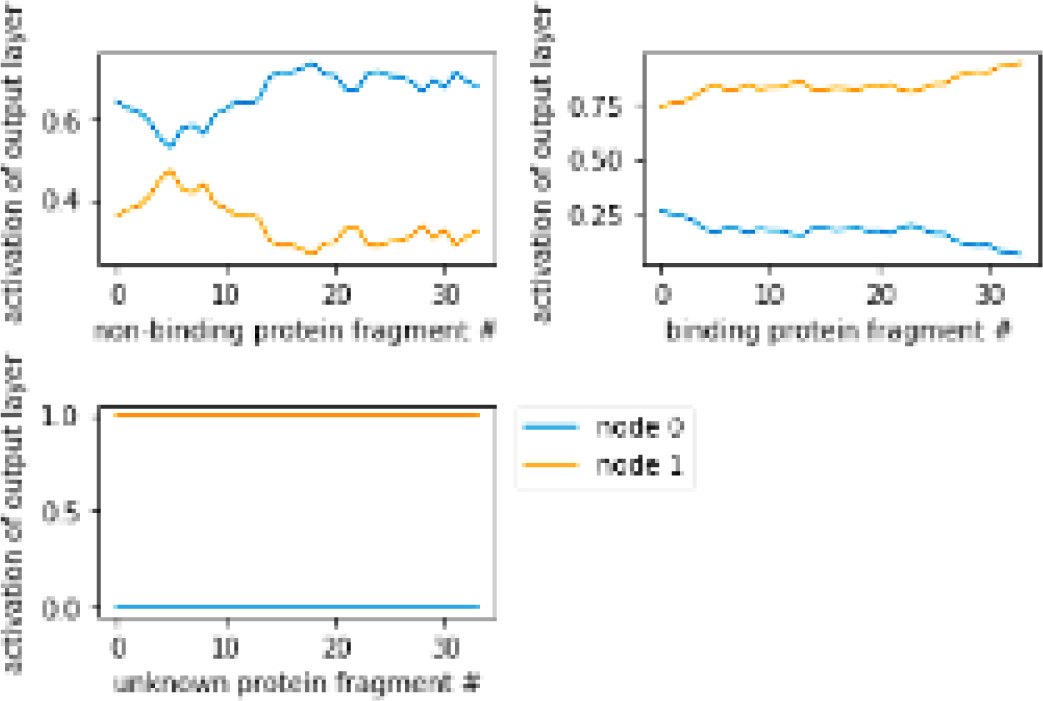
Artemisinin Novel Proteins Binding Classification. Activation of binding node one and not binding node zero for sequence fragments from each of the three proposed proteins to the model for binding classification.

**Fig. 6.**
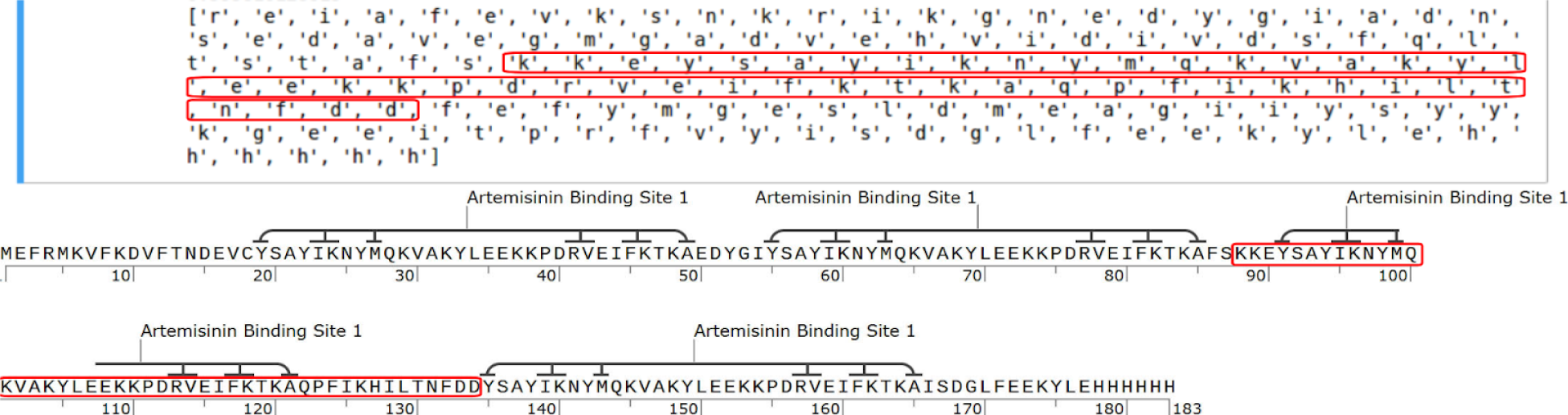
Predicted Binding Sequence. This sequence compares the predicted sequence (top) of specified sequence length for binding by the model to a known binding site found in TCTP (bottom).

**Fig. 7.**
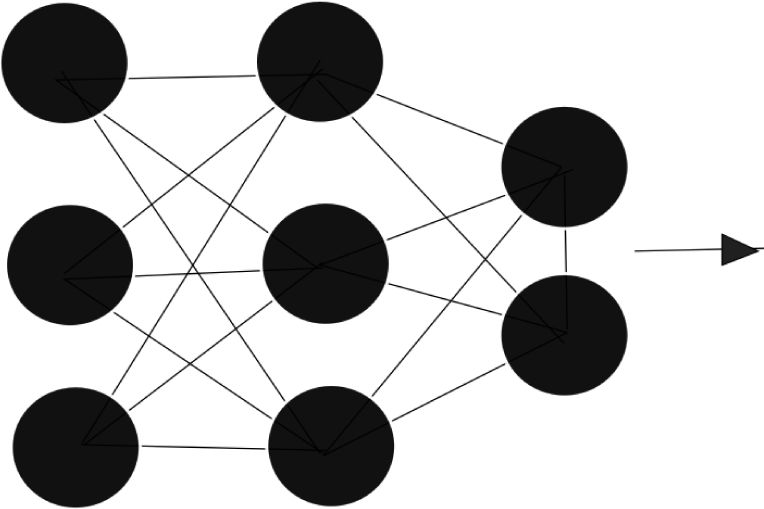
Fully Connected Network. A depiction of a multilayer perceptron, where each node is connected to every node in the adjacent layers.

**Fig. 8.**
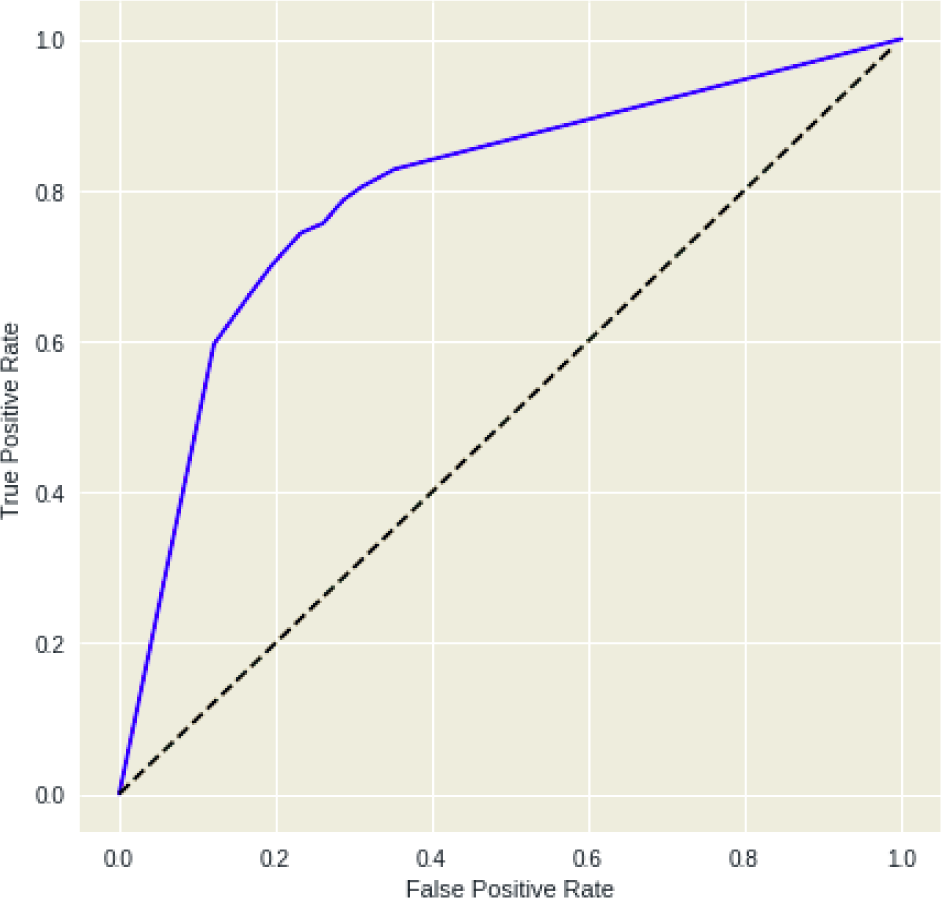
Homeobox ROC Curve. The receiver operator characteristic (ROC) curve of the homeobox model when using string length 50. AUC = 0.80.

**Fig. 9.**
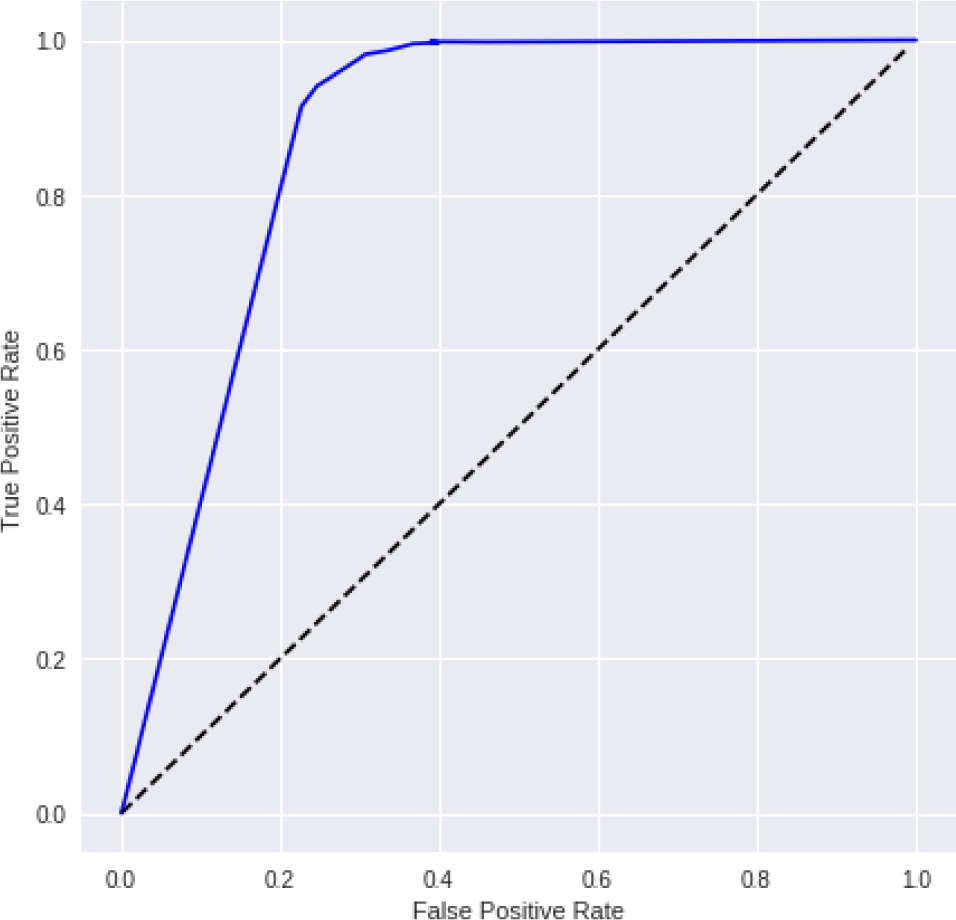
Artemisinin ROC Curve. The receiver operator characteristic (ROC) curve of the Artemisinin model when using string length 50. AUC = 0.87.

## Methods

The architecture proposed used three fully connected LSTM layers built with tensorflow library (Fig. 7). The first two layers comprise 300 nodes taking primary sequence input and previous layer task output, respectively, while the third layer of two nodes informs final decision, based on hidden layer output. The models require unambiguous data pertaining to a certain category and data not in said category. Primary sequence in FASTA amino acid format and classification group were the only inputs given to the model. A dictionary was created by a command code which converted single letter identifiers of amino acid to unique numbers and then back to letters when necessary. Each protein was viewed by the model with a ‘sliding window’ technique, allowing the user to specify sequence input length of each individual protein to be viewed by the model at a each instance. Sequence input length by the ‘sliding window’ technique is also highly useful to manipulate in different types of datasets for different probing purposes, as seen in the two datasets presented. Three useful phazes can be identified during the computational process: training/validation, testing, and analysis for both datasets. In the first phase, the model is trained with 80% of the dataset, 20% for validation. The testing mode is the second phase in which novel (synthetic or arbitrarily composed) sequences are presented to the trained model to predict which category the entire protein belongs to. The third phase can be termed analysis as the model shows the composition of the sequence most related to the specified class (ie. binding domain). Since the models are end-to-end, the analysis is determined by the trained model in a fashion that finds the most conserved feature of the dataset.

### Homeodomain Model

Data used in this model was imported directly from Uniprot’s data-warehouse for proteins containing homeodomain sequences among homosapiens, specified by keyword ‘homeobox’ and organism keyword ‘homo-sapiens’, pulling all associated proteins in the database related to such keyword. Homeodomain was chosen as the key word to establish class from Uniprot because the consensus sequence illustrated in literature is regarded as a highly conserved 60 amino acid long sequence [2]. Again, as data was imported, only primary sequence in FASTA form was retained for training which class the protein belonged to (homeobox containing or not). The proteins in the ‘not’ dataset were imported similarly using keyword ‘NOT-keyword’. This technique pulls a random sample of proteins from the database that are not in the homeobox dataset, creating unique datasets for each class. A command code was utilized to determine the number of proteins to be pulled from Uniprot and used as data. In this case 2000 proteins were used for the dataset in each class. Since the theoretically accepted length of the homeodomain consensus sequence is 60 amino acids long, the sequence length window to be tested was set in a range of 60-100 to ensure the whole sequence would be more likely to be viewed in large chunks rather than smaller individual parts (similar to biological processes). The network tested sequence length in loop increments of five. If a homeodomain is predicted by the network to be present, node one (consensus containing) will have a value between zero and one that is higher than node zero (not consensus containing). During the analysis phase, the model was queried for what sequence composition activated node one the most with a variable sequence length, providing the predicted homeodomain sequence.

### Artemisinin Model

Data for computation in this model was hand sourced from NCBI for proteins known from literature to bind and those known from literature not to bind to the target drug. In this case, Artemisinin was arbitrarily chosen as the target. While 125 binding proteins were found and 130 non-binding proteins were found, the model only used 125 randomly selected from each different class, in order to balance the training set. Since binding mechanisms are mostly unknown and highly variable for the Artemisinin target drug, a range was created from 3-150 amino acid sequence length to be tested. The lower limit of 3 was chosen based on our assumption, derived from the literature, that binding sites are most likely most likely to be at least three amino acids long while the upper limit of 150 was based on computational limits on testing time, which increases exponentially with sequence length. When the network predicted the portion of a protein in the dataset that was likely to bind, a value from zero to one was giving for node zero (non-binding) and node one (binding). Computing differences of nodes to each datapoint in the dataset provides the analysis output of predicted binding motif (as in the homeodomain analysis phase). The least different component shows the inferred compositional property: binding domain.

## Discussion

Overall, the reported results indicated successful use of deep neural networks for protein class prediction and elucidation of functional components. The same deep learning architecture, based on Long Short-Term Memory networks, was successful in inferring two properties from primary sequences: binding predictability of a target drug and consensus sequence detection. While both applications utilized protein’s primary sequence as the dataset and completed the same two tasks successfully: classification and feature selection, it is the responsibility of the scientist to determine the way in which that feature corresponds to the class. Such methodology could even elucidate unknown compositional frameworks in new and old protein datasets. Since previously proposed models are still computationally time consuming, even when running on accelerated hardware, the ability of a model to assess large datasets from cloud sources is novel and highly useful [4,5,10]. Specifically, the efficiency of the proposed LSTM here is valuable, due to its accuracy, expediency, variability, and above all, its ability to utilize a single source of input data: FASTA. Further applications of these types of computation models include integrating various diverse types of datasets for a variety of scientific inquiry. Any type of primary sequence dataset should be able to be assessed by the model as long as general categories and a relationship between them and the data-points can be established. Efficiency may be determined by the abundance of data in the set and the training ability of the network, especially if more than one category will be evaluated. The nature of this computational approach implies that the more data in the set, the increased likelihood of accuracy due to the networks increased ability for training. Further testing will need to be done on the limits of the hardware, such as computational time for larger ‘sliding window’ snippets. It should also be noted that the ability to directly connect to a data source like Uniprot provides the opportunity to eliminate large amounts of pre-processing time, previously used to organize large databases into readable formats as well as hand-annotate features for such data. This work is the first of many to test and employ the model to demonstrate its efficiency and flexibility in a new application, such as proteomics using only primary sequence to predict function. We expect and have foreseen the integration of LSTM-based techniques inomics research to greatly reduce time, cost, and increase accuracy in scientific pipelines such as biology, chemistry, and many other fields that generate large amounts of data that inherently hold information for said research application.

## Data Access

Current datasets can be viewed by querying Uniprot.org for the specific search keywords designated in the Methods section of this work. Original time of Uniprot data access was November 2017. Artemisinin dataset can be found in the appendix section two. The code repository of the models’ architecture can be found in appendix section three.

## Acknowledgments

The authors of this text would like to acknowledge Dr. Nwadiuto Esiobu and Douglas Holmes for suggestion of Artemisinin topic and genetic engineering input, David Dunleavy and the FAU Owlgem team for their input and support, and Nvidia for funding of the GPU hardware used in this research.

## Appendix

### Section 2: Supplementary Data

#### Artemisinin Binding Dataset

https://github.com/MedBios/LSTM-target-bind-/blob/master/BindProteins.txt

#### Artemisinin Non-binding Dataset

https://github.com/MedBios/LSTM-target-bind-/blob/master/NoBindProteins.txt

### Section 3: Code Repository

https://github.com/MedBios/LSTM-target-bind-/tree/master

